# Chronic Physical and Vicarious Psychosocial Stress Alter Fentanyl Consumption and Nucleus Accumbens Rho GTPases in Male and Female C57BL/6 Mice

**DOI:** 10.1101/2021.11.22.469613

**Authors:** Daniela Franco, Andreas B Wulff, Mary Kay Lobo, Megan E Fox

## Abstract

Chronic stress can increase the risk of developing a substance use disorder in vulnerable individuals. Numerous models have been developed to probe the underlying neurobiological mechanisms, however most prior work has been restricted to male rodents, conducted only in rats, or introduces physical injury that can complicate opioid studies. Here we sought to establish how chronic psychosocial stress influences fentanyl consumption in male and female C57BL/6 mice. We used chronic social defeat stress (CSDS), or the modified vicarious chronic witness defeat stress (CWDS), and used social interaction to stratify mice as stress-susceptible or resilient. We then subjected mice to a 15 day fentanyl drinking paradigm in the home cage that consisted of alternating forced and choice periods with increasing fentanyl concentrations. Male mice susceptible to either CWDS or CSDS consumed more fentanyl relative to unstressed mice, and exhibited increased preference for fentanyl. CWDS-susceptible female mice did not differ from unstressed mice during the forced periods, but showed increased preference for fentanyl. We also found decreased expression of nucleus accumbens Rho GTPases in male, but not female mice following stress and fentanyl drinking. We also compare fentanyl drinking behavior in mice that had free access to plain water throughout. Our results indicate that stress-sensitized fentanyl consumption is dependent on both sex and behavioral outcomes to stress.

## Introduction

Repeated and severe stress has long been associated with the emergence of psychiatric disorders including substance use disorders(Sinha, 2008; Hollon et al., 2015; Sapolsky, 2015; Newman et al., 2018; Wemm and Sinha, 2019). Opioid use disorder (OUD) is of particular concern as rates of opioid abuse have skyrocketed, especially in North America (GBD 2016 Alcohol and Drug Use Collaborators, 2018). Accompanying the increase in OUD is an alarming number of overdose deaths and decreased average lifespan (Dowell et al., 2017). Synthetic opioids are a major contributor to increased death rates, and in 2020, caused ~75% of drug overdose deaths in the United States (Ahmad; Skolnick, 2021).

A wealth of literature has sought to uncover how stress drives increased susceptibility to substance use, and the neurobiological underpinnings of comorbid stress and substance use disorders (For recent reviews see: (Newman et al., 2018; Calarco and Lobo, 2020). Psychosocial stress is of particular interest as it is thought to more closely mimic human stressors. However, the effects of psychosocial stress on substance abuse are often divergent and depend on the drug, stress duration, and species (reviewed in (Neisewander et al., 2012). For example in rats, brief social stress increases cocaine self-administration and motivation for cocaine (Haney et al., 1995; Tidey and Miczek, 1997; Covington and Miczek, 2001; Burke and Miczek, 2015), while prolonged social stress reduces cocaine self-administration (Miczek et al., 2011). In mice, prolonged social stress either promotes or suppresses cocaine self-administration (Yap and Miczek, 2007; Han et al., 2015, 2017; Arena et al., 2019) but we recently showed this depends on social housing conditions (Engeln et al., 2021). There are comparatively fewer studies on psychosocial stress and opioid self-administration. In rats, social stress does not influence heroin self-administration (Cruz et al., 2011), while in mice it is associated with increased morphine preference (Cooper et al., 2017).

To investigate how stress influences vulnerability to synthetic opioid use, we adapted an oral fentanyl paradigm in male rats (Shaham et al., 1992) to male and female mice. In the Shaham model, rats were subjected to daily immobilization stress and presented with fentanyl in the homecage drinking water for 4 days (“forced consumption”). Then, rats experienced alternating periods of a choice between fentanyl and water, and additional forced consumption periods. Over time and as concentrations increased, stressed rats increased their fentanyl preference relative to unstressed rats. We sought to replicate this finding in male and female C57BL/6 mice using a similar alternating forced/choice protocol with increasing concentrations. We chose the C57BL/6 mouse since many transgenic tools are developed in this strain (Gong et al., 2007). Like (Cooper et al., 2017), we used the standardized chronic social defeat stress (CSDS) procedure. (Berton et al., 2006; Krishnan et al., 2007; Golden et al., 2011; Fox et al., 2020b, 2020a). CSDS produces anhedonia in the majority of mice (~60%, termed “stress-susceptible”), along with a decrease in motivated behaviors such as social interaction (Krishnan et al., 2007; Der-Avakian et al., 2014; Heshmati and Russo, 2015; Fox and Lobo, 2019). The other ~40% are termed “stress-resilient” and do not display deficits in motivated behavior.

When studying stress and OUD vulnerability, it is also paramount to account for pain-related confounds (Dib and Duclaux, 1982) due the analgesic effects of opioids. Thus, to eliminate pain associated with physical injury (Cooper et al., 2017), we also used chronic witness defeat stress (CWDS), in which mice witness the social defeat of a C57BL/6 mouse and experience “vicarious” social stress (Warren et al., 2013, 2014; Iñiguez et al., 2018). Like CSDS, CWDS produces susceptible and resilient cohorts marked by decreased or unaltered social interaction and motivation, respectively. Since anhedonia and reduced motivation are associated with substance use and dependence (Hatzigiakoumis et al., 2011), we included stress-susceptibility as a factor in our analysis. Finally, we examined how exposure to stress and fentanyl alters expression of dendritic complexity molecules in the Nucleus Accumbens (NAc), a region that is sensitive to both drugs and stress (Hollon et al., 2015; Newman et al., 2018; Calarco and Lobo, 2020). Here we show fentanyl preference after stress is dependent on sex and stress-phenotype, and is associated with altered expression of cytoskeleton-related molecules in the NAc.

## Materials and Methods

### Experimental subjects

All experiments were approved by the Institutional Animal Care and Use Committee at the University of Maryland School of Medicine (UMSOM) and performed in accordance with NIH guidelines for the use of laboratory animals. Mice were given food and water *ad libitum* and housed in the UMSOM vivarium on a 12:12 h light: dark cycle. Experimental mice were 8-9 week old male and female C57BL/6 mice bred at UMSOM. Male CD-1 retired breeders (Charles River, > 4 months) were used as the aggressors for CSDS/CWDS. Mice were randomly assigned to control or stressed groups. The **forced/choice cohort** (see below) included n=12 unstressed males, 12 unstressed females, 23 CSDS males, 19 CWDS males, and 20 CWDS females. The **choice cohort** included n=11 unstressed males, 13 unstressed females, 31 CSDS males, 17 CWDS males, and 15 CWDS females. We excluded 1 unstressed male mouse from this experiment due to abnormally low social interaction behavior.

### Chronic Social Stress

Social stress was performed as in our previous work (Fox et al., 2020a, 2020b; Engeln et al., 2021; Morais-Silva et al., 2021), by using the modifications described in (Warren et al., 2013; Iñiguez et al., 2018) for vicarious resident-intruder stress. In chronic social defeat stress (CSDS), a male mouse (intruder) is physically defeated by an aggressive CD-1 (resident) for 10 min in a hamster cage containing woodchip bedding and a perforated divider. In chronic witness defeat stress (CWDS), a male or female mouse is housed on the opposite side of the perforated divider and allowed to witness the agonistic resident-intruder interactions for 10 min. The CSDS mouse is then housed opposite the resident, and the CWDS mouse is removed and housed opposite to a new CD-1 resident. Following 24h of sensory interaction, the CSDS mouse is defeated by a new CD-1 resident while a different CWDS mouse witnesses the agonistic interaction. This process repeats for 10 days for 10 novel CD1-CSDS-CWDS pairings. Unstressed control mice are pair-housed across perforated dividers in cages containing woodchip bedding with a sex-matched conspecific for 10 days. Immediately following the last stressor, both stressed and unstressed mice are housed individually in woodchip bedding cages.

### Social Interaction Testing

Twenty-four hours after the last stressor, mice were tested for stress-susceptibility in a 3-chamber social preference test (Morais-Silva et al., 2021). Mice were placed in an arena (60 x 40 cm, white walls and floor) divided into 3 chambers (20 x 40 cm) by perforated clear acrylic dividers. The two outer chambers contain wire mesh cups, while the central chamber is empty. The experimental mouse is placed in the central chamber of the arena with two empty wire mesh cups and allowed to explore for 5 min. Then the experimental mouse is allowed to explore the arena for an additional 5 min, this time with unfamiliar sex-matched adult conspecific in one of the wire mesh cups. The amount of time spent in the chamber containing the cups (empty or novel mouse) is measured with video tracking software (TopScan Lite, CleverSys, Reston, VA, USA), and used to determine stress-phenotype. Mice spending <170s in the mouse-paired chamber were deemed “susceptible,” and >170s were “resilient.”

### Homecage fentanyl administration

Following social interaction testing, mice were weighed then pair-housed across a perforated divider with a sex and stress-phenotype matched conspecific in a woodchip bedding cage. Each mouse was provided two 50 mL conical tubes with rubber stoppers and ballpoint sipper tubes (Ancare, Bellmore, NY, USA). All tubes were weighed daily to determine liquid consumption, and the volume consumed was normalized to individual mouse weights.

For the first 4 days (“**forced epoch 1**”), both tubes contained 5 μg/mL fentanyl citrate dissolved in tap water (Cayman # 22659). On day 5, the solution in each mouse’s preferred tube was replaced with plain tap water, and the solution in the least-preferred tube was replaced with 10 μg/mL fentanyl (“**choice 1**”). On days 6-9 (“**forced epoch 2**”), both tubes contained 10 μg/mL fentanyl. On day 10, the preferred tube was replaced with plain water, and the least-preferred tube was replaced with 15 μg/mL fentanyl (“**choice 2**”). On days 11-14 (“**forced epoch 3**”), both tubes contained 15μg/mL fentanyl. On day 15, the preferred tube was replaced with plain water (“**choice 3**”). On day 16, fentanyl solutions were replaced with water and mice were housed individually. Mice were then reassessed for social interaction behavior on day 18. Fentanyl preference was calculated as a percent of total liquid intake.

### RNA isolation

Four 14-gauge NAc tissue punches per mouse were collected 24 hours after the last social interaction and stored at −80°C until processing. RNA was extracted as described previously(Calarco et al., 2021) with Trizol (Invitrogen; #15596018) and the EZNA MicroElute Total RNA kit (Omega BioTek, Norcross, GA; #R6831-01) with a DNase step (Qiagen, Germantown, MD; #79254). RNA concentration and quality were determined with a Nanodrop 1000 spectrophotometer (Thermo Scientific). 400 ng of complementary DNA (cDNA) was synthesized using the reverse transcriptase iScript complementary DNA synthesis kit (Bio-Rad, Hercules, CA; # 1708891), then diluted to a concentration of 2 ng/μL. Relative mRNA expression changes were measured by quantitative PCR using Perfecta SYBR Green FastMix (Quantabio, Beverly, MA; #95072) with a Bio-Rad CFX384 qPCR system. Primer sets are available in **Table 1**. Fold change expression was calculated using the 2^-ddCt^ method with *Gapdh* as the reference gene. Data were normalized to the respective unstressed male or female controls.

### Statistics

All statistical analysis was conducted in GraphPad Prism (version 9, San Diego, California). For CWDS mice, we used 2-way ANOVA with sex and stress-phenotype as factors and Sidak’s post-hoc tests. For CSDS mice, we used 1-way ANOVA with Sidak’s post-hoc, or ANOVA with Welch’s correction and Dunnet’s T3 comparison when standard deviations were significantly different between groups. For within-subject measurements, we used 2-way repeated-measures (RM) ANOVA with Sidak’s post-hoc. For gene expression analysis, we used 2-way ANOVA in CWDS mice, and unpaired t-test for CSDS mice. To compare social interaction between the **forced/choice cohort** and the **choice cohort**, we used 3-way ANOVA using sex, stress-phenotype, and future-group as factors. We used Pearson’s correlation to examine relationships between social interaction and fentanyl preference.

## Results

Numerous studies demonstrate psychosocial stress exerts differential effects on drug-abuse-like behavior (Neisewander et al., 2012; Newman et al., 2018). Previously, we showed individual stress-response impacts cocaine self-administration (Engeln et al., 2021). To test the hypothesis that susceptibility to psychosocial stress increases homecage opioid consumption, we subjected mice to chronic witness defeat (CWDS) or social defeat stress (CSDS) (Timeline and schematic in **Fig 1A**). We divided mice into subgroups based on their behavioral response to stress (Krishnan et al., 2007). Mice that displayed interaction times like unstressed mice were termed “stress-resilient” (>170s), and those that spent less time (<170s) were termed “stress-susceptible” (Example heat maps in **Fig 1B**. CWDS interaction in **Fig1C**: 2-way ANOVA, stress-phenotype F_2,57_ =20.35, *p*<0.0001; unstressed vs CWDS-susceptible and CWDS-susceptible vs CWDS-resilient, Sidak’s post-hoc both p<0.0001. CSDS interaction in **Fig1D**: Welch’s ANOVA, W_2, 16.51_ =21.86, *p*<0.0001)

**Figure 1.**
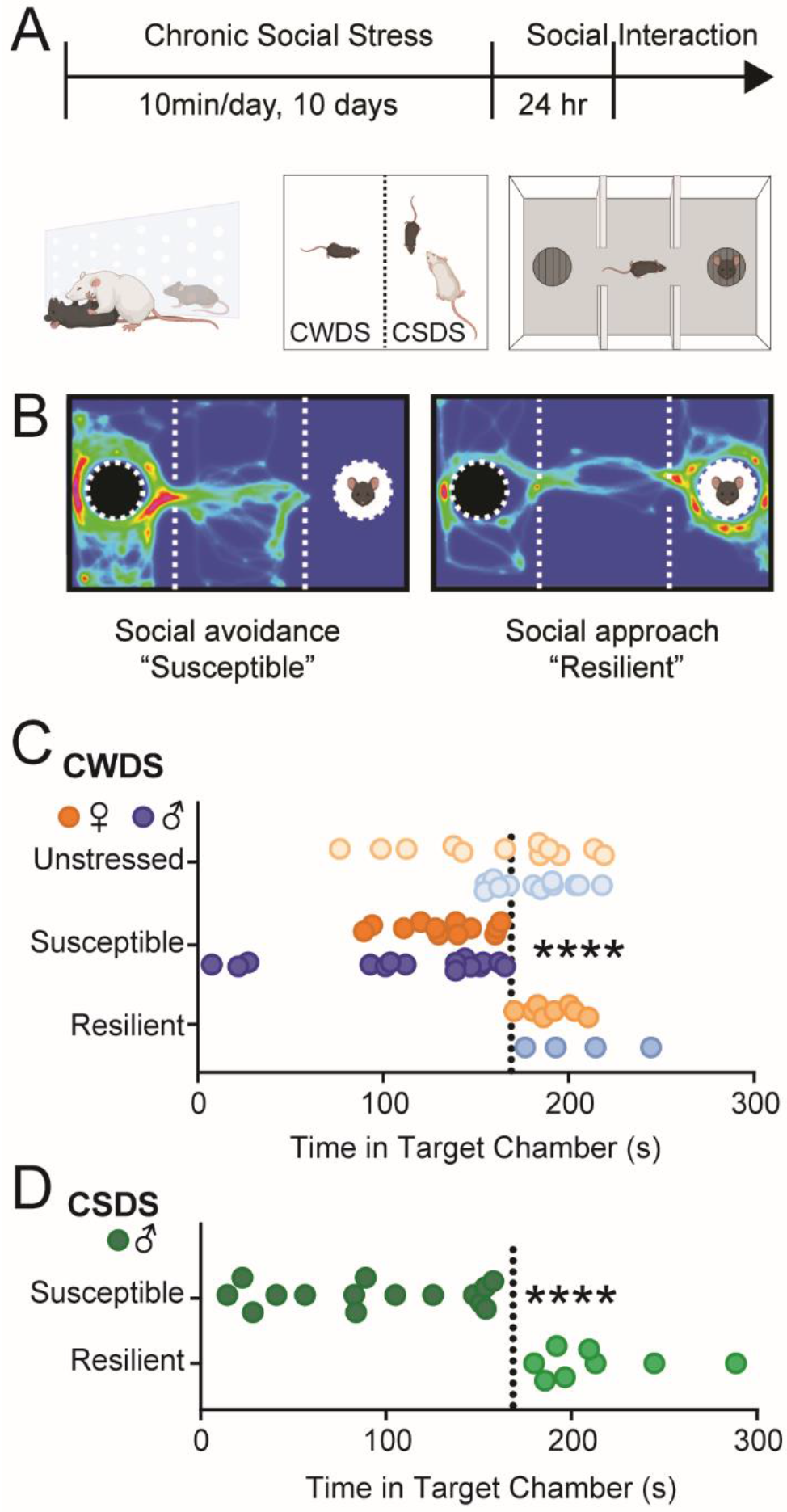
(A) Timeline for chronic social stress. Mice were physically defeated by an aggressive CD-1 resident (CSDS) or witnessed the physical defeat across a perforated divider (CWDS) for 10 days. 24hr after the last stressor, mice were assessed for stress-susceptibility in a 3-chamber social interaction apparatus. (B) Example heatmaps showing susceptible mice spend more time in the non-social compartment containing an empty cup, whereas resilient mice spend more time in the social-compartment containing a novel sex-matched conspecific. (C) Time spent in the social-compartment in unstressed, susceptible, and resilient CWDS male and female mice. Each circle represents an individual mouse. Orange circles are female, blue circles are male. (D) Time spent in the social-compartment containing a novel mouse. Each circle represents an individual mouse. CSDS mice are male only.

### Chronic social stress increases early forced fentanyl consumption in male mice

Following social interaction testing, mice were pair-housed across a perforated divider with a sex and stress-phenotype matched conspecific. Each mouse was provided two tubes containing five μg/mL fentanyl in tap water for 4 days (“**forced epoch 1**”, timeline in **Fig 2**). We compared fentanyl consumption (μg/g bodyweight) during **forced epoch 1** and found a significant effect of stress phenotype (2-way ANOVA; F_2,57_=4.91, *p*=0.011; **Fig3A-B**) and a sex x stress-phenotype interaction in CWDS mice (F_2,57_=3.819, *p*=0.0278). Sidak’s post-hoc tests revealed a sex difference, with unstressed female mice consuming more fentanyl than unstressed males (*p*=0.041, **Fig3B**). We found that CWDS males consume more relative to unstressed males (unstressed vs CWDS-susceptible: *p*=0.012; vs CWDS-resilient: *p*=0.036), however CWDS female mice did not consume more fentanyl relative to unstressed females (*p*>0.05; **Fig3B**). We performed similar comparisons in CSDS mice and found increased fentanyl consumption relative to unstressed male mice (ANOVA, F_2,32_=18.75, *p*<0.0001; **Fig3C-D**). Sidak’s post-hoc tests revealed this increase was dependent on stress-phenotype, with CSDS-susceptible consuming the most fentanyl (unstressed vs CSDS-resilient: *p*=0.094; unstressed vs CSDS-susceptible: p<0.0001; CSDS-susceptible vs CSDS-resilient: *p*=0.014).

**Figure 2.**
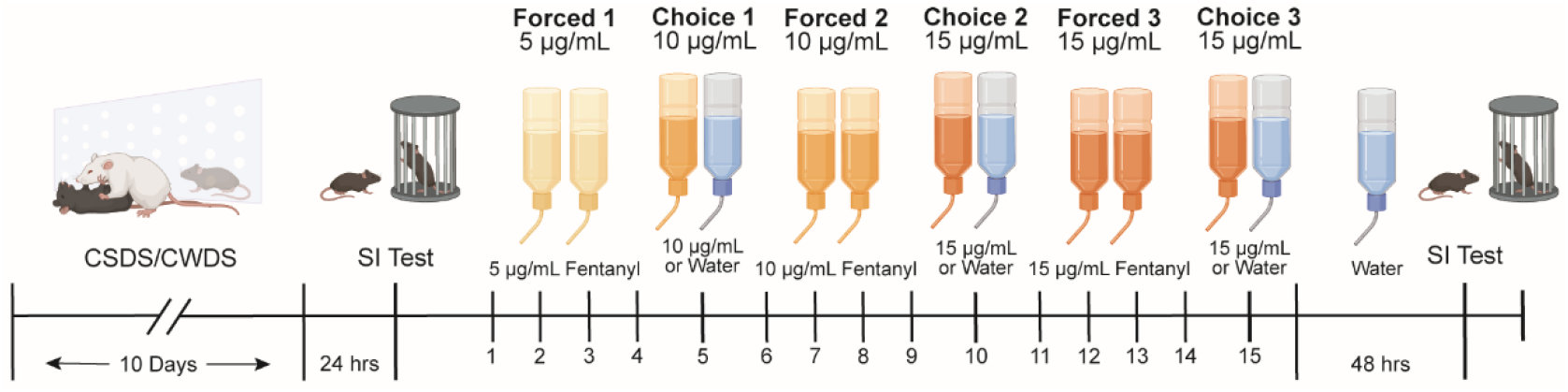
Experimental timeline for forced/choice fentanyl drinking paradigm. Mice underwent 10 days of CSDS or CWDS before completing a social interaction test. Following social interaction testing, mice were organized into behavioral subgroups (susceptible vs. resilient) before starting the forced/choice fentanyl drinking paradigm. Mice experienced 3 alternating periods of forced fentanyl access and 3 periods of non-forced fentanyl access. A second social interaction test was completed ~56 hours after the last day of fentanyl access.

**Figure 3.**
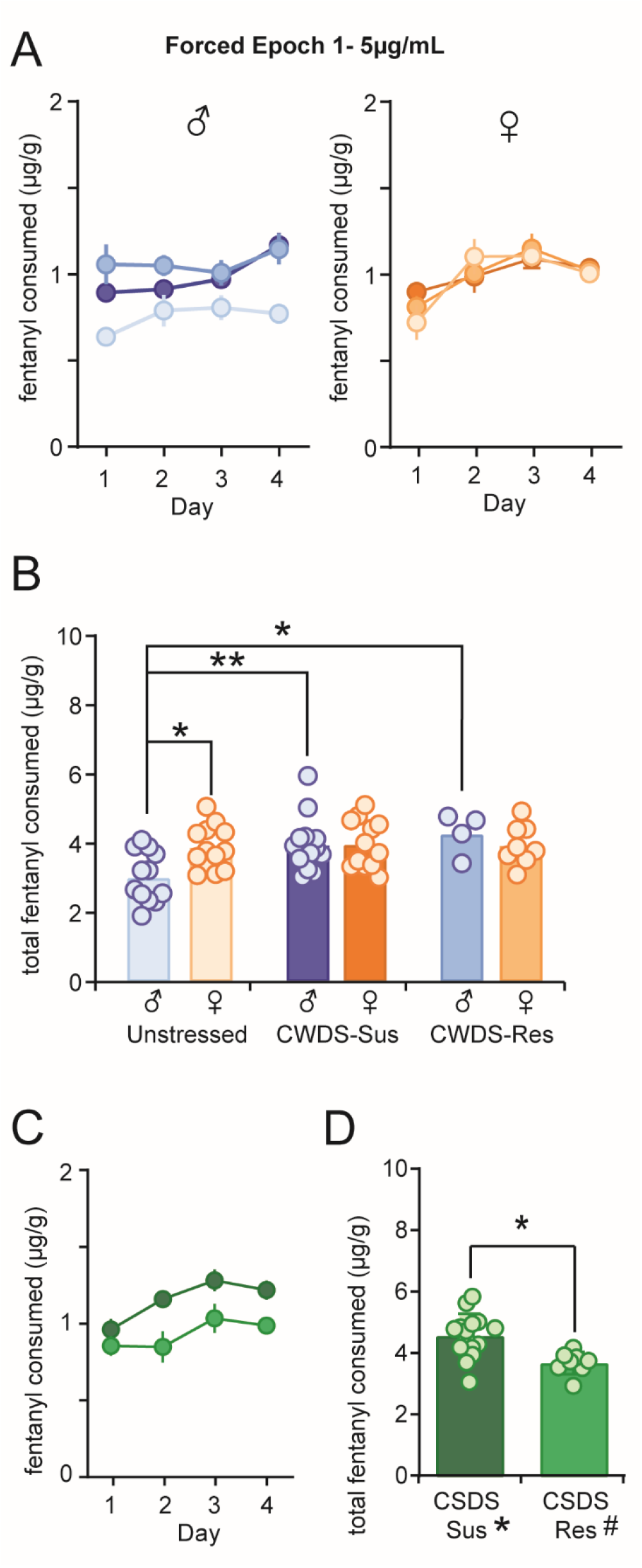
Fentanyl consumption (μg/g) in Forced Epoch 1 (5μg/mL) for unstressed, susceptible, and resilient CWDS and CSDS mice. (A) Average ± SEM fentanyl consumed (μg/g) across days 1-4 for male and female unstressed, CWDS susceptible, and resilient groups. (B) Total fentanyl consumed (μg/g) during Forced Epoch 1. Each circle represents an individual mouse (unstressed male vs unstressed female, \**p* = 0.041; unstressed male vs CWDS-susceptible male, \*\**p* = 0.012; unstressed male vs CWDS-resilient, \**p* = 0.036). (C) Average ± SEM fentanyl consumed (μg/g) across days 1-4 for CSDS susceptible and resilient mice. (D) Total fentanyl consumed (μg/g) during Forced Epoch 1. Each circle represents an individual mouse (unstressed male vs CSDS-susceptible, \**p* < 0.0001; unstressed male vs CSDS-resilient, #*p* = 0.094; CSDS-susceptible vs CSDS-resilient, \**p* = 0.014).

### Stress-susceptible mice exhibit increased early fentanyl preference

On day 5, we replaced the solution in the mouse’s preferred tube with plain tap water, and their least-preferred tube with 10 μg/mL fentanyl (“**Choice 1**”). When we examined fentanyl preference, we found a main effect of stress-phenotype (F_2,58_=3.99, *p*=0.024; **Fig4A**). Sidak’s post-hoc showed CWDS-susceptible, but not CWDS-resilient mice have greater %fentanyl preference relative to unstressed mice (*p*=0.041). We found similar increased μg/g fentanyl consumption during **Choice 1** in CWDS-susceptible, but not CWDS-resilient mice (2-way ANOVA: stress-phenotype F_2,57_=7.51, *p*=0.001; Sidak’s post-hoc *p*=0.001; **Fig 4B**). Physically stressed CSDS mice did not exhibit significantly greater %fentanyl preference relative to unstressed males (ANOVA F_2,32_=1.44, *p*=0.251; **Fig4C**). However, when we examined μg/g fentanyl consumption, we found CSDS-susceptible mice consumed more fentanyl relative to unstressed males (**Fig4D**, Welch’s ANOVA W2,14.02=3.49, *p*=0.05; Dunnet’s T3 comparison: CSDS-susceptible vs unstressed male *p*=0.049).

**Figure 4.**
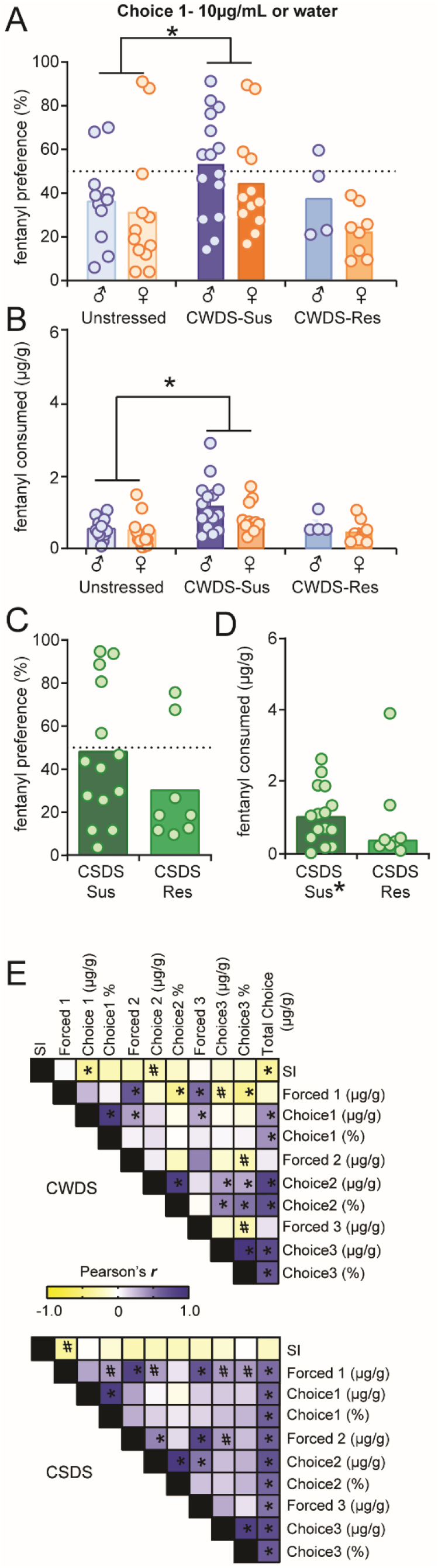
Fentanyl consumption and preference for Choice 1 (10μg/mL or water). (A) Fentanyl preference (%) for Choice 1 (10μg/mL or water) for unstressed, susceptible, and resilient male and female CWDS mice. Each circle represents an individual mouse (unstressed vs CWDS-susceptible, \**p* = 0.041). (B) Fentanyl consumed (μg/g) during Choice 1 (10μg/mL or water) for unstressed, susceptible, and resilient male and female CWDS mice (unstressed vs CWDS-susceptible, \**p* = 0.001). (C) Fentanyl preference (%) for Choice 1 (10μg/mL or water) for CSDS mice. (D) Fentanyl consumed (μg/g) during Choice 1 (10μg/mL or water) for CSDS mice (unstressed male vs CSDS-susceptible, \**p* = 0.049). (E) Pearson correlation matrix representing paired correlations between stress-phenotype (measured by social interaction, SI) and fentanyl preference. Negative correlations are in yellow; positive correlations are in blue. *, p<0.05. #, p<0.09.

### Stress-susceptibility drives early fentanyl consumption in CWDS mice

Given that stress-susceptible mice consumed more fentanyl during **Forced Epoch 1** and **Choice 1**, we next sought to strengthen the relationship between stress-phenotype and fentanyl preference. We performed Pearson’s correlations using social interaction data from individual mice. In CWDS mice, we found a negative correlation between social interaction and μg/g fentanyl consumed during **Choice 1**, indicating that the less time spent interacting with a novel conspecific (i.e. more stress-susceptible), the more fentanyl the CWDS mouse consumed during **Choice 1** (Pearson’s *r*=-0.382, *p*=0.026; **Fig4E, top**). By contrast, stress-susceptibility was not significantly correlated with μg/g fentanyl consumed during **Choice 1** in CSDS mice (*p*=0.94, **Fig4E, bottom**). Of note, there was no significant relationship between social interaction and μg/g fentanyl consumption in unstressed mice (not shown, *p*=0.18).

### Stress-susceptible male mice continue to consume more fentanyl in **Forced Epoch 2**

On day 6, we provided mice with two tubes containing 10 μg/mL fentanyl for 4 days (“**Forced Epoch 2**”). We compared fentanyl consumption during **Forced Epoch 2**, and found a significant effect of stress-phenotype (F_2,57_=8.52, *p*=0.0006) and a sex x stress-phenotype interaction (F_2,57_=6.74, *p*=0.002, **Fig5A-B**). Like **Forced Epoch 1**, male CWDS-susceptible and CWDS-resilient consumed more μg/g fentanyl than unstressed male mice (*p*<0.0001 and *p*=0.010, respectively), while female CWDS mice consumed amounts equivalent to unstressed females. Again, stress also increased fentanyl consumption in CSDS mice dependent on stress-phenotype, with CSDS-susceptible consuming the most μg/g in **Forced Epoch 2** (ANOVA: F_2,32_=10.84, *p*=0.0003. Sidak’s post-hoc: CSDS-susceptible vs unstressed, *p*=0.0002; CSDS-resilient vs unstressed, *p*=0.187; **Fig 5C-D**).

**Figure 5.**
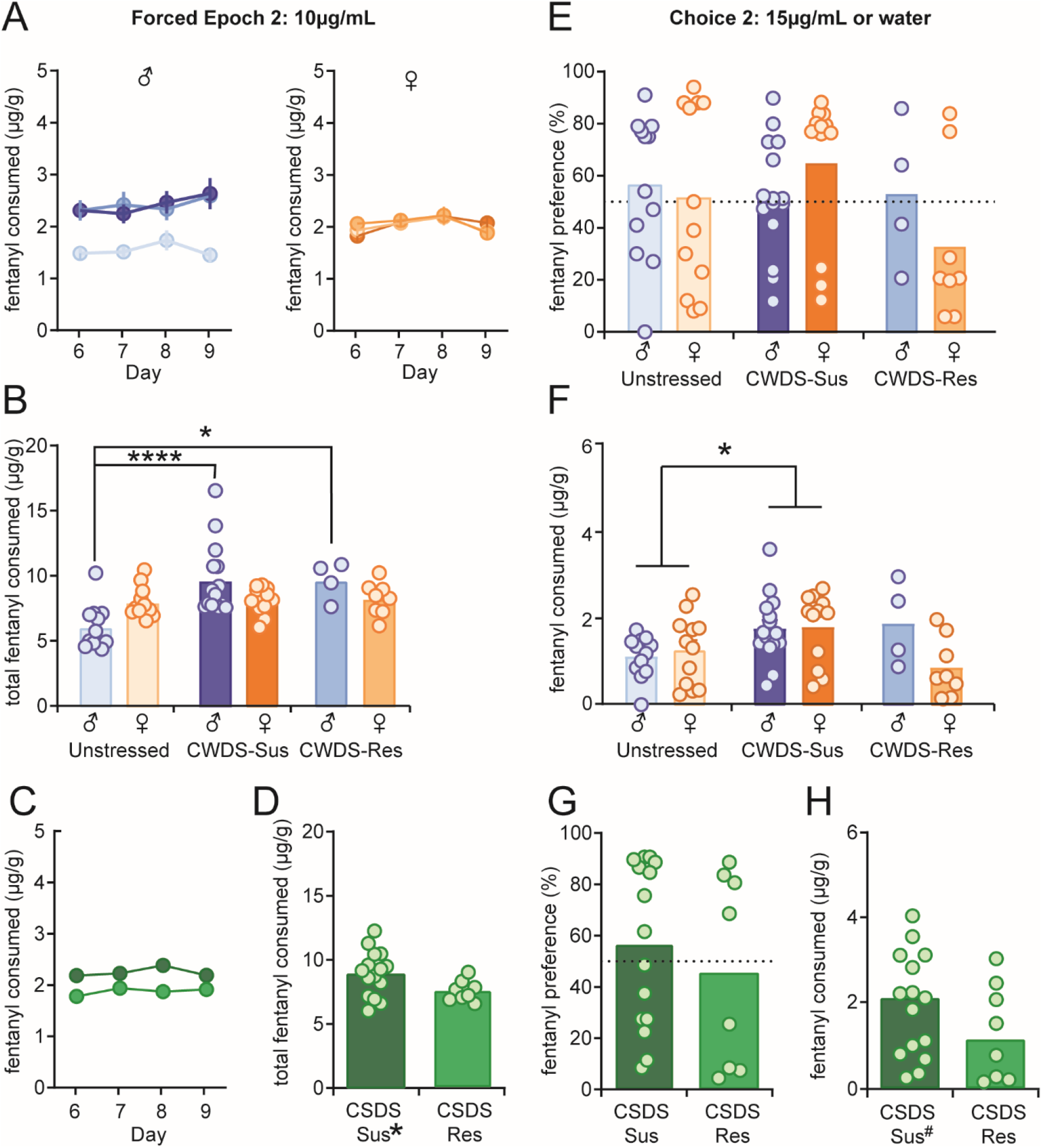
Fentanyl consumption (μg/g) in Forced Epoch 2 (10μg/mL) and Choice 2 (15μg/mL) for unstressed, susceptible, and resilient CWDS and CSDS mice. (A) Average ± SEM fentanyl consumed (μg/g) across days 69 for male and female unstressed and CWDS susceptible and resilient groups, respectively. (B) Total fentanyl consumed (μg/g) during Forced Epoch 2. Each circle represents an individual mouse (unstressed male vs CWDS-susceptible male, \*\*\*\**p* < 0.0001; unstressed male vs CWDS-resilient male, \**p* = 0.010). (C) Average ± SEM fentanyl consumed (μg/g) across days 6-9 for CSDS susceptible and resilient mice. (D) Total fentanyl consumed (μg/g) during Forced Epoch 2 (unstressed male vs CSDS-susceptible, \**p* = 0.0002; unstressed male vs CSDS-resilient, *p* = 0.187). (E) Fentanyl preference (%) for Choice 2 (15μg/mL) for unstressed, susceptible, and resilient male and female CWDS mice. (F) Fentanyl consumed (μg/g) during Choice 2 (15μg/mL or water) for unstressed, susceptible, and resilient male and female CWDS mice (unstressed vs CWDS-susceptible, \**p* = 0.02). (G) Fentanyl preference (%) for Choice 2 (15μg/mL or water) for CSDS mice. (H) Fentanyl consumed (μg/g) during Choice 2 (15μg/mL or water) for CSDS mice (unstressed male vs CSDS-susceptible, #*p* = 0.0590).

### Stress-susceptibility drives increased fentanyl consumption during **Choice 2** in CWDS mice

On Day 10, we replaced the solution in mice’s least preferred tube with 15 ug/mL fentanyl, and their preferred tube with plain water (“**Choice 2**”). We examined preference for the fentanyl tube and found no significant effect of stress-phenotype or sex (**Fig 5E**). When we examined μg/g consumption however, we found a significant effect of stress-phenotype (2-way ANOVA, F_2,57_=4.10, *p*=0.022) and CWDS-susceptible mice consumed more fentanyl during **Choice 2** than unstressed mice (Sidak’s post-hoc, p=0.02; **Fig5F**). Similar to **Choice 1**, we found a trending negative correlation between stress-susceptibility and μg/g fentanyl consumed during **Choice 2** (Pearson’s *r*=-0.270, *p*=0.097; **Fig4E**). CSDS mice did not exhibit greater %fentanyl preference during **Choice 2** (ANOVA: F_2,32_=0.36 p=0.702; **Fig5G**), but showed a trend towards greater fentanyl consumption relative to unstressed males (Dunnet’s T3 unstressed vs CSDS-susceptible *p*=0.0590; **Fig5H**). Like **Choice 1**, μg/g fentanyl consumption during **Choice 2** was not correlated with stress-susceptibility in CSDS mice (*p*=0.287, **Fig4E**).

### Stress-susceptible male mice continue to consume more fentanyl in **forced epoch 3**

On day 11, we provided mice with two tubes containing 15 μg/mL fentanyl for 4 days (“**forced epoch 3**”). We examined consumption during **forced epoch 3** in CWDS mice, and found a significant effect of stress-phenotype (F_2,57_=5.72, *p*=0.005) and a sex x stress-phenotype interaction (F_2,57_=9.21, *p*=0.0003, **Fig6A-B**). As with **Forced Epoch 1** and **2**, male CWDS-susceptible and CWDS-resilient mice consumed more fentanyl than unstressed male mice (Sidak’s post-hoc, unstressed vs susceptible *p*=0.0001, vs. resilient *p*=0.0002; **Fig 6B**), while female mice showed no differences relative to unstressed controls. Physically stressed CSDS mice also consumed more compared with unstressed males (ANOVA, F_2,32_=12.98, *p*<0.0001; Sidak’s post-hoc unstressed vs susceptible, *p*<0.0001; vs. resilient, *p*=0.092; **Fig6C-D**).

**Figure 6.**
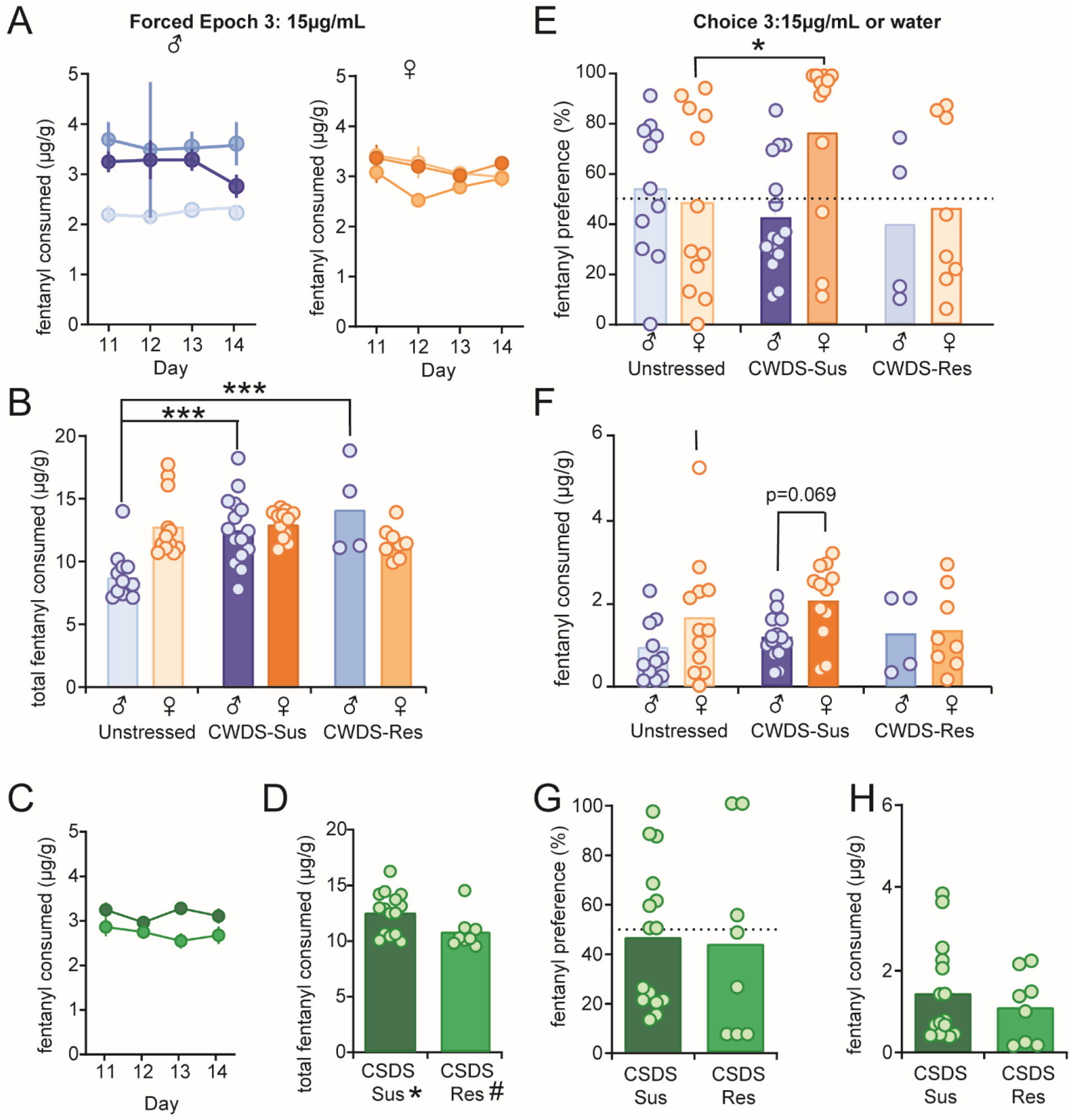
Fentanyl consumption (μg/g) in Forced Epoch 3 (15μg/mL) and Choice 3 (15μg/mL) for unstressed, susceptible, and resilient CWDS and CSDS mice. (A) Average ± SEM fentanyl consumed (μg/g) across days 11-14 for male and female unstressed, CWDS susceptible and CWDS-resilient groups, respectively. (B) Total fentanyl consumed (μg/g) during Forced Epoch 3. Each circle represents an individual mouse (unstressed male vs CWDS-susceptible male, \*\*\**p* = 0.0001; unstressed male vs CWDS-resilient male, \*\*\**p* = 0.0002). (C) Average ± SEM fentanyl consumed (μg/g) across days 11-14 for CSDS susceptible and resilient mice. (D) Total fentanyl consumed (μg/g) during Forced Epoch 3 (unstressed male vs CSDS-susceptible, \**p* < 0.0001; unstressed male vs CSDS-resilient, #*p* = 0.092). (E) Fentanyl preference (%) for Choice 3 (15μg/mL) for unstressed, susceptible, and resilient male and female CWDS mice. \**p*=0.046 (F) Fentanyl consumed (μg/g) during Choice 3 (15μg/mL or water) for unstressed, susceptible, and resilient male and female CWDS mice (CWDS-susceptible male vs CWDS-susceptible female, *p* = 0.069). (G) Fentanyl preference (%) for Choice 3 (15μg/mL or water) for CSDS susceptible and resilient mice. (H) Fentanyl consumed (μg/g) during Choice 3 (15μg/mL or water) for CSDS mice.

### Stress-susceptible female, but not male mice, consume more fentanyl during **Choice 3** while behavior is decoupled from consumption

On day 15, we replaced the mouse’s preferred tube with plain water and assessed preference for the 15μg/mL fentanyl solution (“**Choice 3**”). In CWDS mice, we found a trending stress phenotype x sex interaction (2-way ANOVA, F_2,57_=3.01, *p*=0.057; **Fig6E**). Male CWDS mice exhibited similar fentanyl preferences regardless of stress phenotype. Female CWDS-susceptible mice showed a higher preference for the fentanyl bottle compared with unstressed females (Sidak’s post hoc, *p*=0.046; **Fig 6E**). We examined μg/g fentanyl consumption alongside %fentanyl preference and found a main effect of sex (2-way ANOVA, F_1,57_=4.28, *p*=0.043; **Fig6F**). Female CWDS-susceptible mice consumed more μg/g fentanyl compared with male CWDS-susceptible mice, although this failed to reach statistical significance (Sidak’s post-hoc, male vs female CWDS-susceptible, *p*=0.069; **Fig6F**). By this point in the experiment, stress-susceptibility was no longer correlated with μg/g or %fentanyl preference on **Choice 3** (p=0.372, **Fig4E**). As with **Choice 2**, there was no effect of stress-phenotype on **Choice 3** %fentanyl preference in the CSDS males (ANOVA, F_2,32_=0.43, *p*=0.655), **Choice 3** fentanyl consumption (F_2,32_=0.91, *p*=0.414, **Fig6G-H**), nor a correlation between stress-susceptibility and **Choice 3** consumption (*p*=0.484, **Fig4E**).

### Only stress-susceptible female mice increase fentanyl preference over time

To compare our findings to those in (Shaham et al., 1992), we also examined preference for fentanyl solution as a function of the first and last choice days. Surprisingly, only female CWDS-susceptible mice increased %fentanyl preference on **Choice 3** relative to **Choice 1** (2-way RMANOVA. Time: F_1,57_=6.61, *p*=0.013; Group: F_5, 57_=2.57, *p*=0.036; **Choice 1** vs **3**, Sidak’s post-hoc, *p*=0.036; **Fig 7B**). In contrast, male CWDS mice did not exhibit increased %fentanyl preference increased their **Choice 3** preference relative to **Choice 1,** but maintained consistent %preference throughout (p’s >0.05, **Fig 7A**). Male CSDS mice also did not significantly increase **Choice 3** %fentanyl preference relative to **Choice 1** %fentanyl preference (2-way RMANOVA, Day: F_1,32_=2.46, *p*=0.1266, **Fig 7C**).

**Figure 7.**
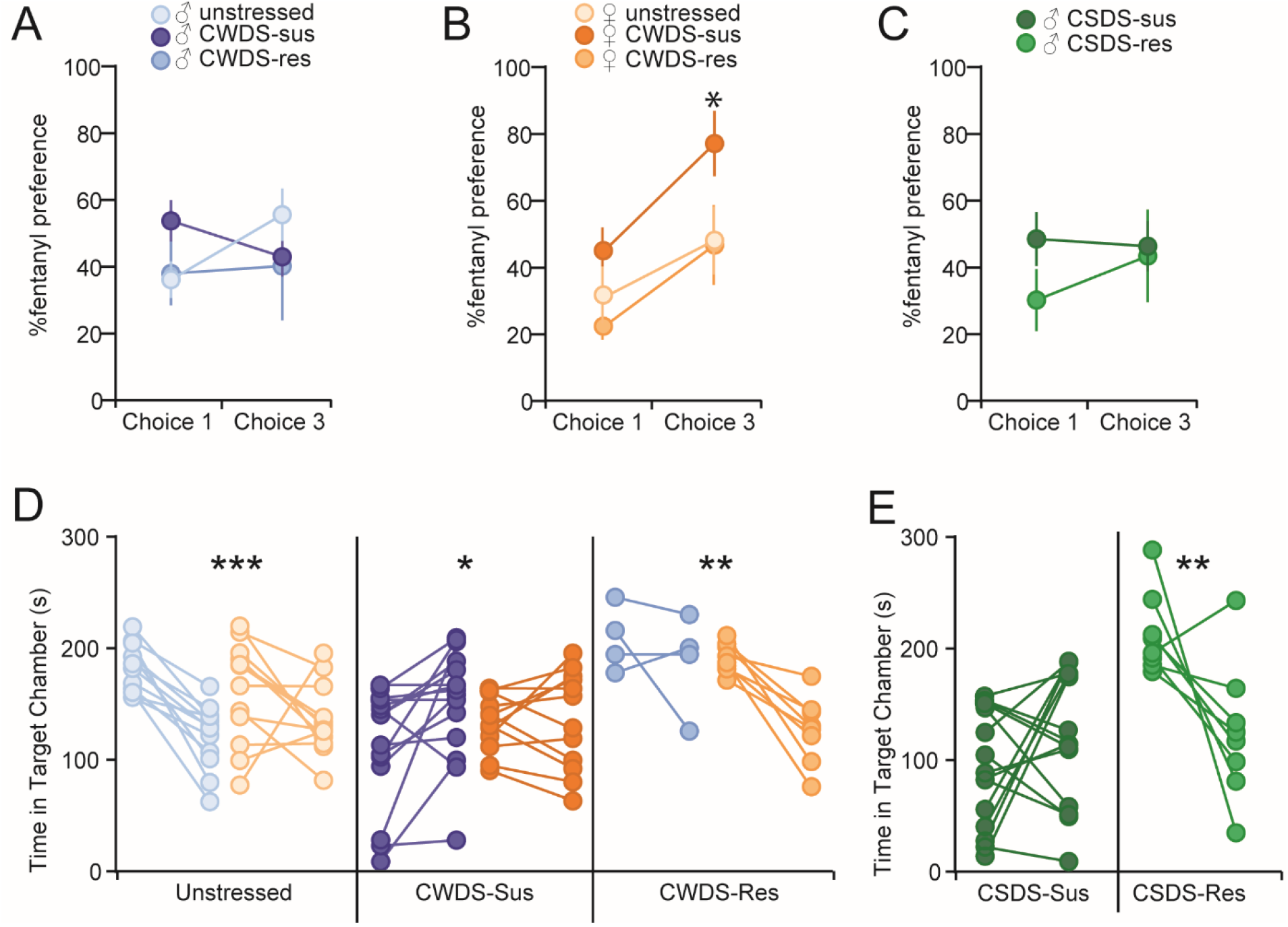
Changes in fentanyl preference and social interaction for unstressed, susceptible, and resilient CWDS and CSDS mice in the Forced fentanyl cohort. (A) Fentanyl preference (%) during Choice 1 and Choice 3 for unstressed, CWDS susceptible, and CWDS resilient male mice. (B) Fentanyl preference (%) during Choice 1 and Choice 3 for unstressed, CWDS susceptible, and CWDS resilient female mice (CWDS-susceptible Choice 1 vs Choice 3, \**p* = 0.036). (C) Fentanyl preference (%) during Choice 1 and Choice 3 for CSDS susceptible and CSDS resilient mice. (D) Time in target chambers (s) during the first social interaction (before fentanyl administration) and second social interaction (after fentanyl administration) for unstressed, susceptible, and resilient CWDS and CSDS mice (unstressed before vs after fentanyl, \*\*\**p* = 0.0002; CWDS-susceptible before vs after fentanyl, \**p* = 0.0465; CWDS-resilient before vs after fentanyl, \*\**p* = 0.0022). (E) Time in target chambers (s) during the first social interaction and second social interaction for CSDS susceptible and resilient mice (CSDS-resilient before vs after fentanyl, \*\**p* = 0.0041).

### Abstinence from homecage fentanyl increases stress-susceptibility dependent on stress-phenotype

Opioid abstinence and withdrawal can promote anhedonia and social interaction deficits(Bai et al., 2014; Becker et al., 2017; Bravo et al., 2020). To test how our stress and fentanyl paradigm influenced social interaction, we next replaced the fentanyl solutions with plain water for 2 days and single housed mice prior to reassessing social interaction behavior. We chose this ~56 hr time point to ensure complete fentanyl clearance (Kalvass et al., 2007) and to avoid acute withdrawal symptoms during testing. We found significant effects of stress-phenotype (2-way RMANOVA, F_2,60_=6.79, *p*=0.032), time (F_1,60_=12.69, *p*=0.0007), and a time x stress-phenotype interaction (F_2,60_=15.44, p<0.0001; **Fig7D**). All unstressed and CWDS-resilient groups showed reduced time interacting with a novel sex-matched conspecific relative to the first social-interaction (Sidak’s post-hoc, before vs after fentanyl: unstressed *p*=0.0002, CWDS-resilient *p*=0.0022, **Fig7D**). CWDS-susceptible showed a nominal increase in social interaction, but remained susceptible (*p*=0.0465). CSDS mice also reduced time interacting with a novel conspecific relative to the first social interaction, but this was only statistically significant in CSDS-resilient mice, possibly indicating a “floor effect” in CSDS-susceptible mice (2-way RMANOVA, time: F_1,32_=10.35, *p*=0.0030; stress-phenotype: F_2,32_=12.90, *p*<0.0001; time x stress-phenotype: F_2,32_=8.93, *p*=0.0008;Sidak’s post-hoc, before vs after fentanyl: CSDS-susceptible *p*=0.3310; CSDS-resilient *p*=0.0041, **Fig 7E**). Together, this indicates fentanyl exposure and abstinence alone is sufficient to generate a stress-like phenotype.

### Chronic stress and fentanyl exposure downregulate dendritic complexity molecules in the Nucleus Accumbens

Both chronic stress and opioid exposure cause structural changes in the nucleus accumbens (NAc) that are thought to drive stress-susceptibility or increased drug intake (Matsubara et al., 1999; Robinson et al., 2002; Spiga et al., 2005; Diana et al., 2006; Christoffel et al., 2011b, 2011a; Pal and Das, 2013; Golden et al., 2013; Guegan et al., 2016; Kobrin et al., 2016; Graziane et al., 2016; Francis et al., 2017; Cahill et al., 2018; Geoffroy et al., 2019; Fox et al., 2020a, 2020b). We thus examined the consequences of our combined stress and fentanyl paradigm on genes associated with dendritic remodeling (Nakayama et al., 2000; Negishi and Katoh, 2005; Newey et al., 2005; Chen and Firestein, 2007). We extracted NAc RNA from a subset of mice, and found decreased expression of Rho GTPases RhoA, Rac1, and Cdc42 in CWDS-male, but not CWDS-female mice. (2-way ANOVA. *RhoA:* Stress, F_1, 27_=9.36, Sex x Stress F_1, 27_=21.65, **Fig 8A**; *Rac1:* Stress F_1, 27_=16.17, Sex x Stress F_1, 27_ =15.70, **Fig 8B**; *Cdc42:* Stress F_1, 27_ =11.68, Sex x Stress F_1, 27_=27.79, **Fig 8C**; all p≤0.005. Sidak’s post-hoc unstressed-male vs CWDS-male, all p<0.0001). We found similar decreased expression of Rho GTPases in CSDS mice (unpaired t-test. *RhoA:* t_13_=5.75, **Fig 8E;** *Rac1:* t_13_=6.34, **Fig 8F;** *Cdc42:* t_13_ =5.14, **Fig 8G;** all p≤0.0002). In contrast, we found no changes in the downstream effector Limk1 (Meng et al., 2003) in CWDS mice of either sex (2-way ANOVA, p>0.05, **Fig 8D**). In CSDS mice we found a trend towards increased Limk1 expression (t_13_=2.06, p=0.060, **Fig 8H**). It is possible the sex and stressor-dependent changes in gene expression are related to sex differences in fentanyl consumption and stress responsivity, which will require further investigation. Nevertheless, we demonstrate that stressed males show downregulation of Rho GTPases. Targeting dendritic remodeling may prove useful for blunting stress-sensitized acquisition of drug taking, and this is worth further investigation (Rigoni et al., 2021).

**Figure 8.**
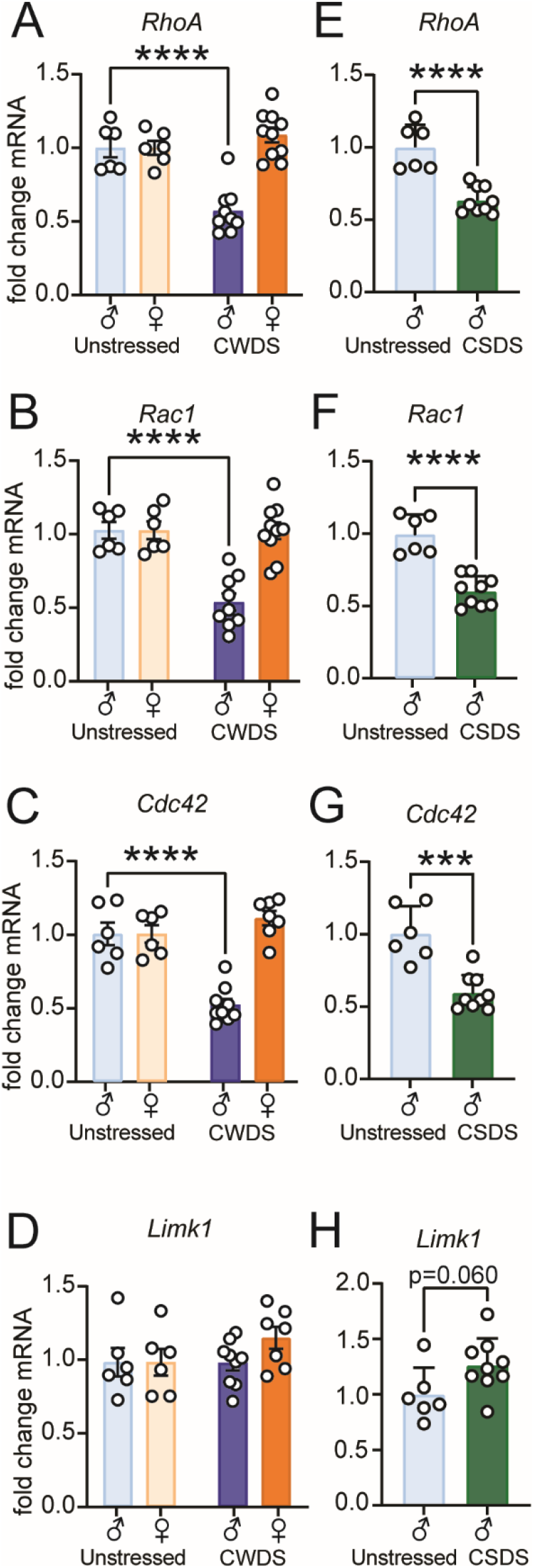
Rho GTPases RhoA, RAc1, and Cdc42 are downregulated in the Nucleus Accumbens after chronic stress and fentanyl exposure in male mice. (A-D) RhoA, Rac1, Cdc42, and Limk1 gene expression in unstressed and CWDS mice, respectively (unstressed male vs CWDS male: *RhoA, Rac1, Cdc42, Limk1, ****p* < 0.0001). (E-H) RhoA, Rac1, Cdc42, and Limk1 gene expression in unstressed and CSDS mice, respectively (unstressed vs CSDS: *RhoA, Rac1, ****p* < 0.0001; unstressed vs CSDS: *Cdc42, ***p* = 0.0002; unstressed vs CSDS: *Limk1, p* = 0.060).

### Female mice exhibit greater fentanyl preference in an all-choice paradigm

An important caveat in this work is that by forcing mice to consume fentanyl, some mice may have increased fentanyl preference due to tolerance or may consume fentanyl to mitigate withdrawal symptoms. We thus repeated the experiment in a separate cohort of mice that were given a choice between fentanyl and water for the entire experiment (“**choice cohort**”, timeline in **Fig 9A**). As with the mice that received forced exposure (“**forced/choice cohort**”), we divided **choice cohort** mice into susceptible and resilient as described above. (For CWDS, 2-way ANOVA, stress-phenotype, F_2,50_ =9.36, *p*=0.0004; **Fig 9B**, For CSDS, Welch’s ANOVA, W2,17.43=30.67, *p*<0.0001; **Fig 9C**). While we had different proportions of susceptible and resilient mice as compared with the **forced/choice cohort**, there was no effect of future-group (3-way ANOVA, F_1,107_ =0.88 *p*=0.3516) or future-group x stress-phenotype (F_2,107_=2.10, *p*=0.13) in CWDS mice. For CSDS mice, there was a future-group x stress-phenotype interaction (2-way ANOVA, F_2,71_ =6.02, p=0.0038) such that CSDS-susceptible mice in the **choice cohort** had greater mean interaction time (Sidak’s post-hoc, **forced/choice cohort** vs **choice cohort**, *p*=0.004). We include the data from this group in the present manuscript, but it is important to note this difference when comparing the two cohorts.

**Figure 9.**
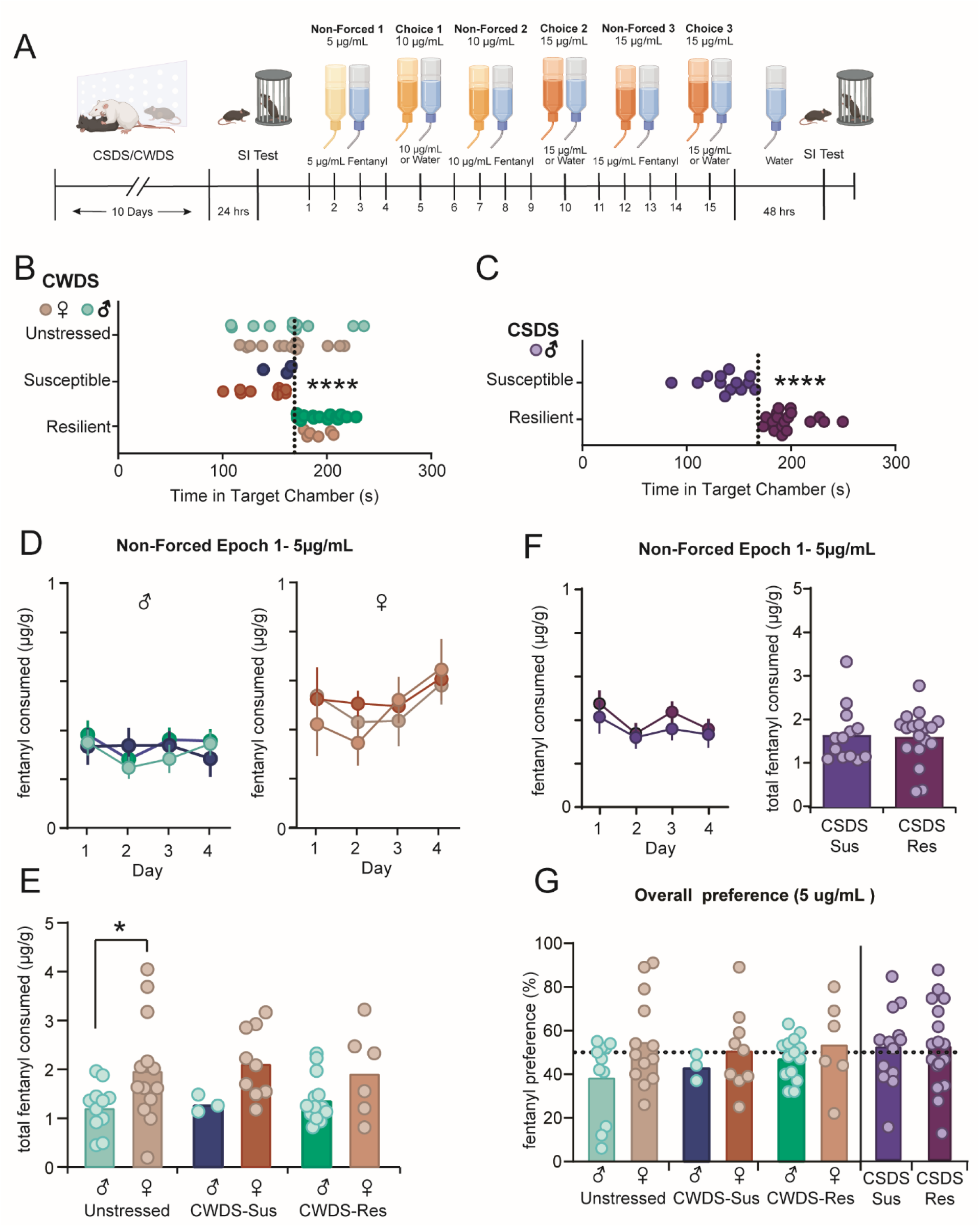
(A) Experimental timeline for non-forced fentanyl drinking paradigm in “choice cohort” mice. Mice underwent 10 days of CSDS or CWDS before completing a social interaction test. Following social interaction testing, mice were organized into behavioral subgroups (susceptible vs. resilient) before starting the non-forced fentanyl drinking paradigm. A second social interaction test was completed ~56 hours after the last day of fentanyl access. (B) Time in target chamber (s) for unstressed, CWDS susceptible, and CWDS resilient male and female mice (\*\*\*\**p* = 0.0004). (C) Time in target chamber (s) for CSDS susceptible and resilient mice (\*\*\*\**p* < 0.0001). (D) Average ± SEM fentanyl consumed (μg/g) during Non-Forced Epoch 1 (5μg/mL or water) across days 1-4 for male and female unstressed, CWDS susceptible and resilient mice, respectively. (E) Total fentanyl consumed (μg/g) during Non-Forced Epoch 1 (5μg/mL or water). Each circle represents an individual animal (unstressed female vs unstressed male, \**p* = 0.048). (F) Average ± SEM fentanyl consumed (μg/g) during Non-Forced Epoch1 (5μg/mL or water) across days 1-4 for CSDS mice (left). Total fentanyl consumed (μg/g) during Non-Forced Epoch 1 (5μg/mL or water) for CSDS susceptible and resilient mice (right). (G) Overall fentanyl preference (%) across days 1-4 (5μg/mL) for unstressed, susceptible, and resilient CWDS and CSDS mice.

Following social interaction, we pair-housed mice with a sex and phenotype-matched mouse. Each mouse was provided with one tube containing 5 μg/mL fentanyl and a second tube containing plain tap water for four days (“**non-forced epoch 1**”). Tube positions were alternated daily to account for side preference. When we examined fentanyl consumption during **non-forced epoch 1**, we found a main effect of sex (2-way ANOVA, F_1, 50_ =9.79, *p*=0.003, **Fig 9D-E**), but no effect of stress. Sidak’s post-hoc revealed unstressed female mice consumed more μg/g fentanyl as compared with unstressed male mice (*p*=0.048). We examined overall %fentanyl preference during **non-forced epoch 1** and found a trending sex effect (2-way ANOVA, F_1, 50_=3.64, *p*=0.06, **Fig 9G**), with unstressed females exhibiting a trend towards greater %fentanyl preference compared with unstressed males (Sidak’s post-hoc, *p*=0.077). CSDS mice in the **choice cohort** did not consume more fentanyl relative to unstressed males (ANOVA F_2,39_=2.02, p=0.147, **Fig 9F**) nor exhibit increased %fentanyl preference (ANOVA, F_2,39_ =1.94, p=0.158, **Fig 9G**).

### Female mice exhibit increased fentanyl consumption in an all-choice paradigm

On day 5, we replaced 5μg/mL fentanyl with 10 μg/mL (**Choice 1**). In CWDS mice, we found a significant stress phenotype x sex interaction (2-way ANOVA, F_2,50_=3.47, *p*=0.039, **Fig 10A**). Sidak’s post-hoc revealed a trend towards increased %fentanyl preference in male CWDS-resilient mice compared with male CWDS-susceptible (Sidak’s post-hoc *p*=0.069). We found a main effect of sex in μg/g consumption during **Choice 1** (2-way ANOVA, F_1, 50_=5.12, *p*=0.028, **Fig 10B**), with CWDS-susceptible females consuming more fentanyl than male CWDS-susceptible (Sidak’s post-hoc *p*=0.028). Despite the group-level differences between CWDS-susceptible and resilient mice, there was no correlation between **Choice 1** %fentanyl preference and social interaction time in CWDS mice (p=0.77, not shown). There were no differences in CSDS mice relative to unstressed males (ANOVA, %fentanyl preference, F_2,39_=0.272, p=0.764, **Fig 10C**; μg/g consumption F_2,39_=1.36, p=0.269, **Fig 10D**).

**Figure 10.**
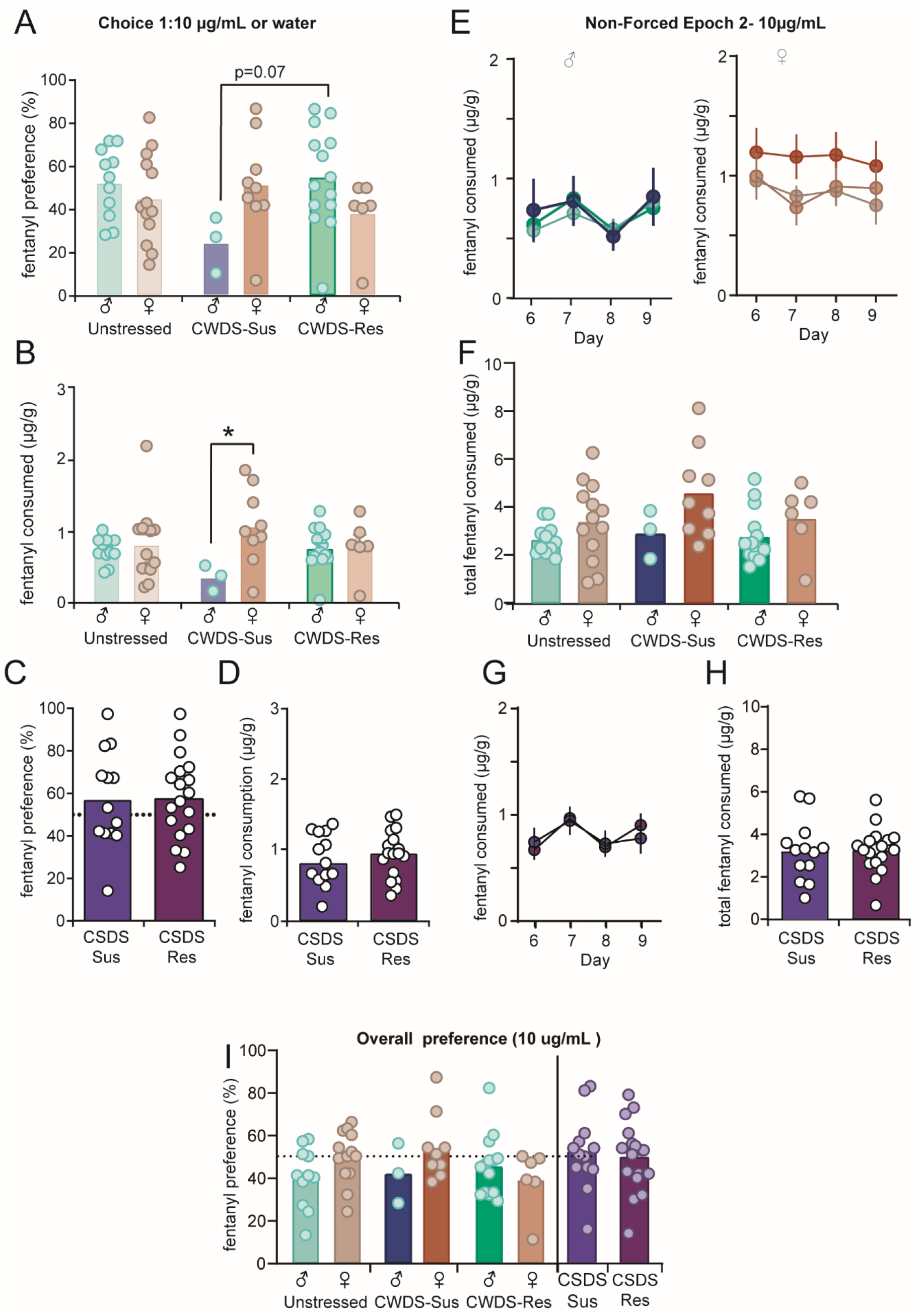
Fentanyl preference (%) and consumption (μg/g) for Choice 1 and Non-Forced Epoch 2 (10μg/mL or water) for unstressed, susceptible, and resilient CWDS and CSDS “choice-cohort” mice. (A) Fentanyl preference (%) for Choice 1 (10μg/mL or water) for male and female unstressed and CWDS susceptible and resilient mice. Each circle represents an individual mouse (CWDS-susceptible male vs CWDS-resilient male, *p* = 0.07). (B) Fentanyl consumed (μg/g) during Choice 1 (10μg/mL or water) for male and female unstressed and CWDS susceptible and resilient mice (CWDS-susceptible male vs CWDS-susceptible female, \**p* = 0.028). (C) Fentanyl preference (%) for Choice 1 (10μg/mL or water) for CSDS mice. (D) Fentanyl consumption (μg/g) during Choice 1 (10μg/mL or water) for CSDS susceptible and resilient mice. (E) Average ± SEM fentanyl consumed during Non-Forced Epoch 2 (10μg/mL or water) across days 6-9 for male and female unstressed and CWDS susceptible and resilient mice. (F) Total fentanyl consumed (μg/g) during Non-Forced Epoch 2 (10μg/mL or water) for male and female unstressed and CWDS susceptible and resilient mice. (G) Average ± SEM fentanyl consumed (μg/g) during Non-Forced Epoch 2 (10μg/mL or water) across days 6-9 for CSDS susceptible and resilient mice. (H) Total fentanyl consumed (μg/g) during Non-Forced Epoch 2 (10μg/mL or water) for CSDS susceptible and resilient mice. (I) Overall fentanyl preference (%) across days 5-9 (10μg/mL) for unstressed, susceptible, and resilient CWDS and CSDS mice.

To mimic the time-course of the **forced/choice cohort** mice, we provided the **choice cohort** with 10 μg/mL fentanyl and water during days 6-9 (**“non-forced epoch 2**”). When we examined fentanyl consumption during **non-forced epoch 2**, we found a main effect of stress (2way ANOVA, F_1, 50_=6.61, *p*=0.013, **Fig 10F**), but no significant differences in post-hoc analysis. We examined the overall %preference for 10μg/mL fentanyl and found no effects of stress or sex in CWDS mice (2-way ANOVA, p>0.05, **Fig 10I**). CSDS mice in the choice cohort did not differ from unstressed mice in μg/g consumption (ANOVA, p=0.342, **Fig 10H**) nor %fentanyl preference (ANOVA, p=0.17, **Fig 10I**).

On day 10, we replaced the 10μg/mL fentanyl with 15 μg/mL (**Choice 2**). In CWDS mice, there was no effect of sex or stress on %fentanyl preference (2-way ANOVA, p>0.05, **Fig 11A**). However, when we examined μg/g consumption, we found a main effect of sex (2-way ANOVA, F_1, 50_=15.29, *p*=0.0003; **Fig 11B**). Female CWDS-susceptible mice consumed more fentanyl compared with male CWDS-susceptible mice (Sidak’s post-hoc, *p*=0.029, **Fig 11B**), however there was no correlation between social interaction and %fentanyl preference (*p*=0.74). There were no differences in CSDS mice (ANOVA, %fentanyl preference:, F_2,39_=0.519, p=0.599, **Fig 11C**; μg/g consumption: F_2,39_=0.335, *p*=0.717, **Fig 11D**).

**Figure 11.**
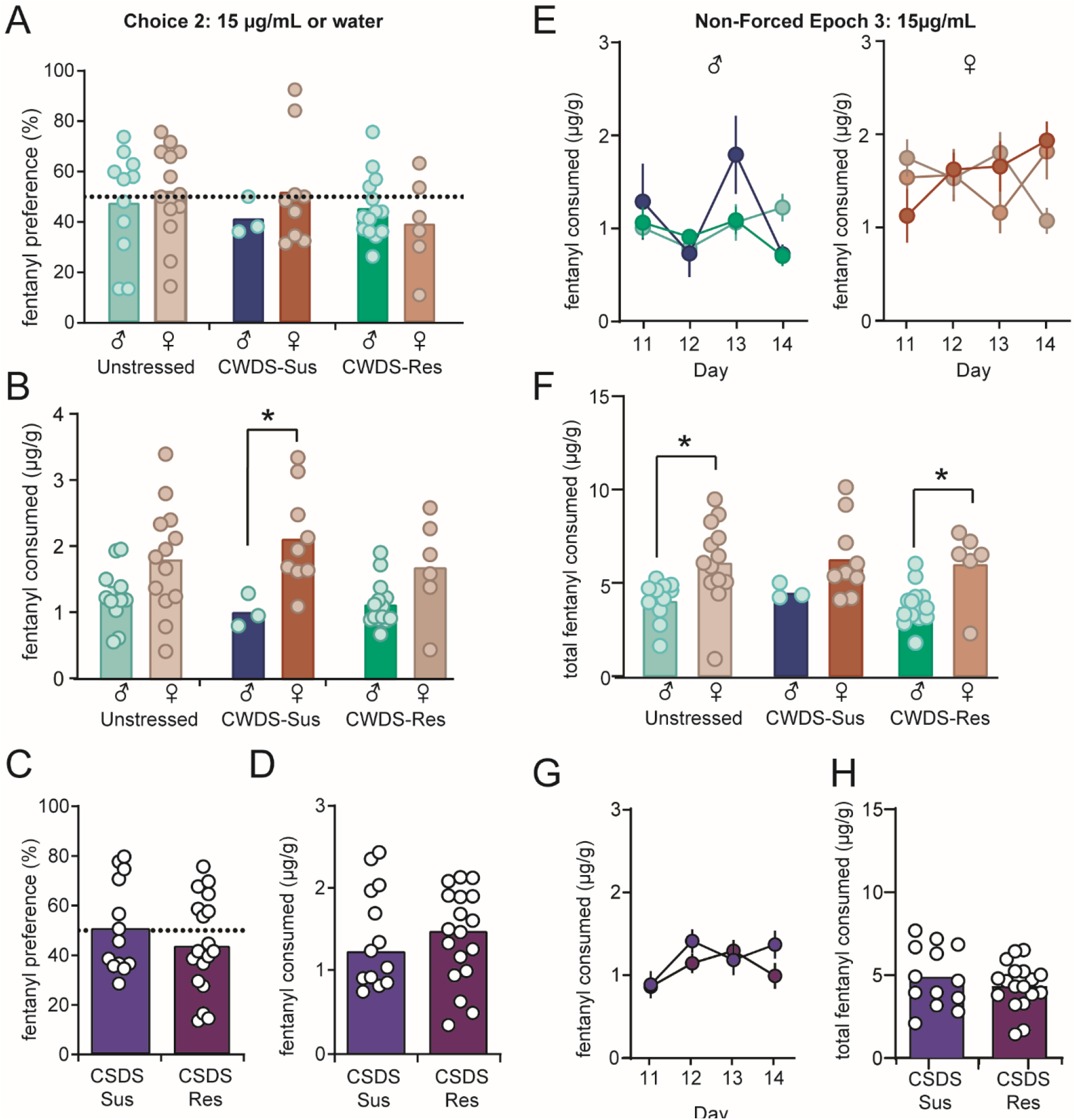
Fentanyl preference (%) and consumption (μg/g) for Choice 2 and Non-Forced Epoch 3 (15μg/mL or water) for unstressed, susceptible, and resilient CWDS and CSDS “choice-cohort” mice. (A) Fentanyl preference (%) for Choice 2 (15μg/mL or water) for male and female unstressed and CWDS susceptible and resilient mice. Each circle represents an individual mouse. (B) Fentanyl consumed (μg/g) during Choice 2 (15μg/mL or water) for male and female unstressed and CWDS susceptible and resilient mice (CWDS-susceptible male vs CWDS-susceptible female, \**p* = 0.029). (C) Fentanyl preference (%) for Choice 2 (15μg/mL or water) for CSDS mice. (D) Fentanyl consumption (μg/g) during Choice 2 (15μg/mL or water) for CSDS susceptible and resilient mice. (E) Average ± SEM fentanyl consumed during Non-Forced Epoch 3 (15μg/mL or water) across days 11-14 for male and female unstressed and CWDS susceptible and resilient mice. (F) Total fentanyl consumed (μg/g) during Non-Forced Epoch 3 (15μg/mL or water) for male and female unstressed and CWDS susceptible and resilient mice (unstressed male vs unstressed female, \**p* = 0.012; CWDS-resilient male vs CWDS-resilient female, \**p* = 0.019). (G) Average ± SEM fentanyl consumed (μg/g) during Non-Forced Epoch 3 (15μg/mL or water) across days 11-14 for CSDS susceptible and resilient mice. (H) Total fentanyl consumed (μg/g) during Non-Forced Epoch 3 (15μg/mL or water) for CSDS susceptible and resilient mice.

To mimic the **forced/choice cohort**, mice were provided with 15 μg/mL fentanyl or water for days 11-14 (“**non**-**forced epoch 3**”). We found a main effect of sex on μg/g consumption in CWDS mice (2-way ANOVA, F_1, 50_ =16.12, *p*=0.0002, **Fig 11E-F**). Sidak’s post-hoc revealed unstressed female mice consumed more fentanyl compared with unstressed males (*p*=0.012, **Fig 11F**), and CWDS-resilient females consumed more compared with CWDS-resilient males (p=0.019, **Fig 11F**). There were no differences in **non-forced epoch 3** consumption in CSDS mice (ANOVA, p=0.379, **Fig 11G-H**). Mice continued to have access to 15 μg/mL and water tubes on day 15 (**Choice 3**). We found a trending sex x stress-phenotype interaction (F_2,50_=2.77, p=0.072; **Fig 12A**) for %fentanyl preference during **Choice 3**. Sidak’s post-hoc revelead CWDS-resilient males had reduced **Choice 3** %fentanyl preference relative to unstressed males (*p*=0.017, **Fig 12A**). We also found a negative correlation between social interaction and **choice 3** preference (Pearson’s r=-0.404, *p*=0.0219), such that more susceptible mice exhibited greater %fentanyl preference. For **Choice 3** μg/g consumption, we found a main effect of sex (2-way ANOVA, F_1, 50_=12.95, *p*=0.0007) with CWDS-susceptible and CWDS-resilient females consuming more compared with their male counterparts (Sidak’s post-hoc, *p*=0.049, *p*=0.038, respectively, **Fig 12B**). There were no differences in CSDS mice (ANOVA, %preference: *p*=0.212, **Fig 12C**; μg/g: p=0.822, **Fig 12D**). We also examined the overall %preference for 15μg/mL fentanyl in CWDS and CSDS mice and found no differences (**Fig12E**).

**Figure 12.**
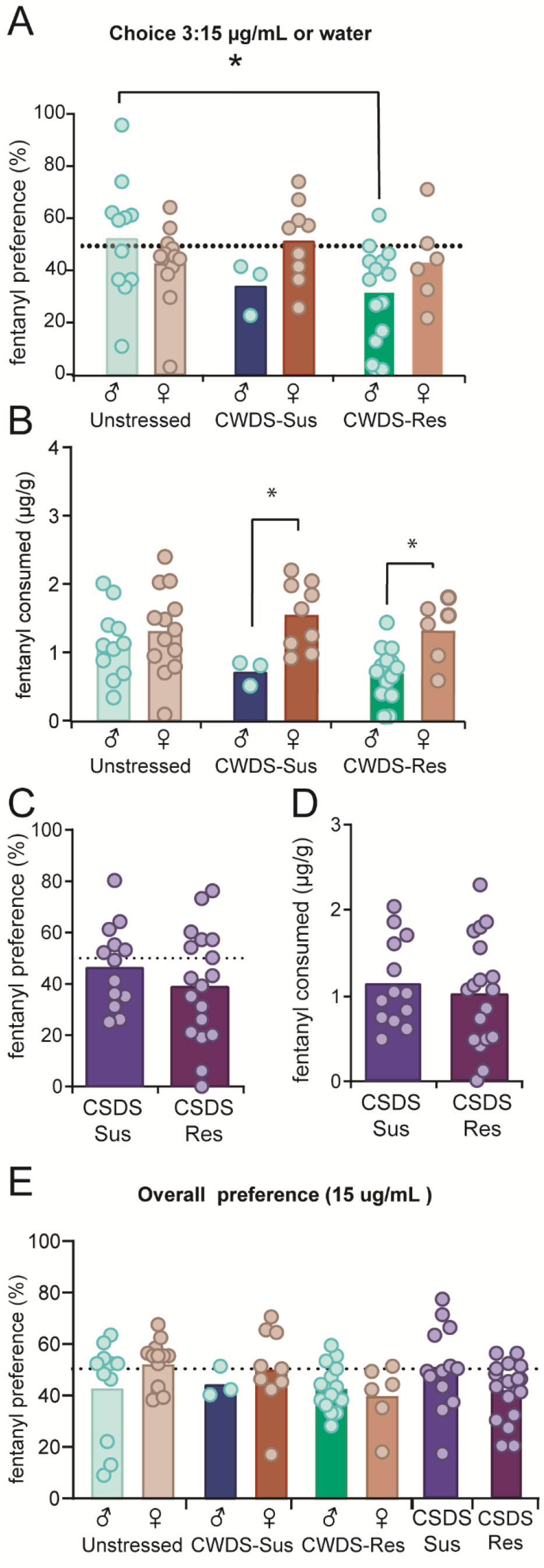
Fentanyl preference (%) and consumption (μg/g) for Choice 3 (15μg/mL or water) for unstressed, susceptible, and resilient CWDS and CSDS “choice-cohort” mice. (A) Fentanyl preference (%) for Choice 3 (15μg/mL or water) for male and female unstressed and CWDS susceptible and resilient mice. Each circle represents an individual mouse (unstressed male vs CWDS-resilient male, \**p* = 0.017). (B) Fentanyl consumed (μg/g) during Choice 3 (15μg/mL or water) for male and female unstressed and CWDS susceptible and resilient mice (CWDS-susceptible male vs CWDS-susceptible female, \**p* = 0.049; CWDS-resilient male vs CWDS-resilient female, \**p* = 0.038). (C) Fentanyl preference (%) for Choice 3 (15μg/mL or water) for CSDS susceptible and resilient mice. (D) Fentanyl consumption (μg/g) during Choice 3 (15μg/mL or water) for CSDS susceptible and resilient mice. (E) Overall fentanyl preference (%) across days 11-14 (15μg/mL) for unstressed, susceptible, and resilient CWDS and CSDS mice.

### In an “all-choice”paradigm, resilient males decrease fentanyl preference over time, but still show social withdrawal after fentanyl

As with the **forced/choice cohort**, we examined %fentanyl preference as a function of **Choice 1** vs **3**. There was a significant time x group interaction for CWDS mice (2-way RMANOVA, F_5,50_=2.43, *p*=0.0477, **Fig 13A-B**), with male CWDS-resilient mice significantly decreasing their preference between the two time points (Sidak’s post-hoc, *p*=0.0037). Female mice maintained %fentanyl preference. There was a main effect of time in CSDS mice (2-way RMANOVA, F_1,39_=4.93, *p*=0.0322 **Fig 13C**), with CSDS-resilient mice decreasing %fentanyl preference between the two time points (Sidak’s post-hoc, *p*=0.016).

**Figure 13.**
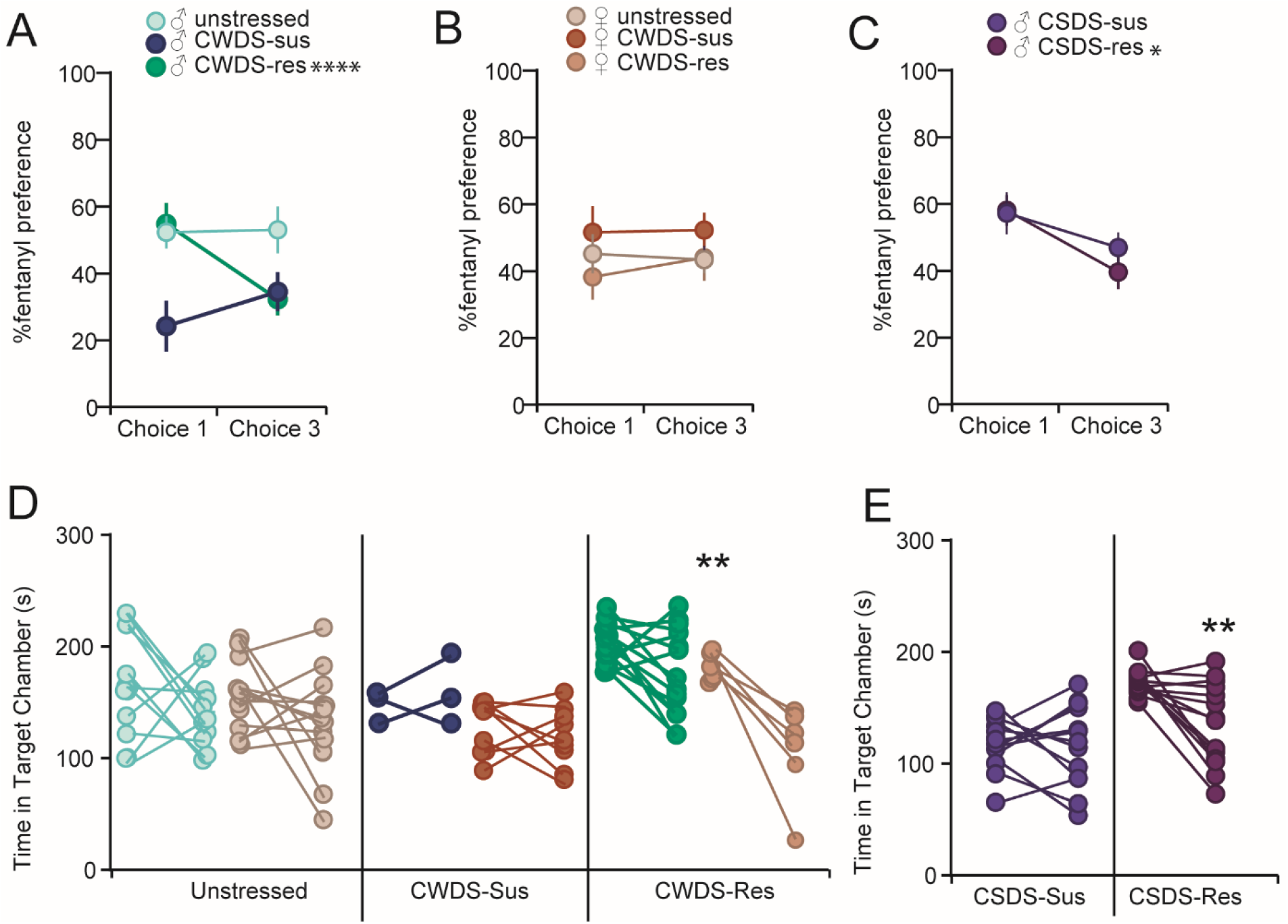
Changes in fentanyl preference and social interaction for unstressed, susceptible, and resilient CWDS and CSDS mice in the “choice cohort.” (A) Fentanyl preference (%) during Choice 1 and Choice 3 for unstressed, CWDS susceptible, and CWDS resilient male mice. (CWDS-resilient male, \*\*\*\**p* = 0.0037). (B) Fentanyl preference (%) during Choice 1 and Choice 3 for unstressed, CWDS susceptible, and CWDS resilient female mice. (C) Fentanyl preference (%) during Choice 1 and Choice 3 for CSDS susceptible and CSDS resilient mice. (CSDS-resilient, \**p* = 0.016). (D) Time in target chambers (s) during the first social interaction (before fentanyl administration) and second social interaction (after fentanyl administration) for unstressed, susceptible, and resilient CWDS and CSDS mice (CWDS-resilient before vs after fentanyl, \*\**p* = 0.0029). (E) Time in target chambers (s) during the first social interaction and second social interaction for CSDS susceptible and resilient mice (CSDS-resilient before vs after fentanyl, \*\**p* = 0.002).

Similar to the **forced/choice cohort**, we retested mice for social interaction after 2 days of plain water. We found a main effect of time (2-way RM-ANOVA, F_1,53_=8.67, *p*=0.005) and stress-phenotype (F_2,53_=8.82, *p*=0.0005; **Fig 13D**). Sidak’s post-hoc showed that only CWDS-resilient mice decreased time spent interacting with a novel conspecific (*p*=0.0029). Similarly in the CSDS mice, only CSDS-resilient mice decreased social interaction time (2-way RM-ANOVA, time F_1,36_=8.36, *p*=0.006; stress-phenotype F_2,36_=8.28, *p*=0.001, Sidak’s post-hoc, p=0.002; **Fig13E)**. Given the lower amount of fentanyl consumption in the unstressed mice, this may reflect either a dose-dependent effect of fentanyl exposure and abstinence on social withdrawal, or fentanyl abstinence is a great enough insult to tip resilient mice towards susceptibility.

## Discussion

Here we show both physical (CSDS) and vicarious (CWDS) psychosocial stressors impact homecage opioid consumption and preference in a stress and sex-dependent manner. First, susceptibility to either CWDS or CSDS increases forced fentanyl consumption in male mice regardless of fentanyl concentration. Second, male and female CWDS mice show increased fentanyl consumption during the first two choice periods, and greater stress-susceptibility is associated with greater fentanyl preference. However only female CWDS-susceptible mice show increased fentanyl preference during the final choice period. While physical CSDS-susceptibility is associated with increased fentanyl consumption during the first choice period, this effect does not persist over the subsequent choices and is decoupled from social interaction behavior. When mice are instead given a choice between fentanyl and water through the entire experiment, only female CWDS-susceptible mice show increased fentanyl consumption. Finally, we show CWDS and CSDS-susceptible males have downregulated Rho GTPases in the NAc.

The main goal of this study was to determine if psychosocial stress in mice produced similar effects to restraint stress in rats on fentanyl preference, using the alternating periods of forced and choice consumption described by (Shaham et al., 1992). Consistent with this prior work, our stressed mice consumed more fentanyl than unstressed mice during the choice epochs. Vicarious stress was sufficient to increase drug-taking, similar to (Ramsey and Van Ree, 1993; Cooper et al., 2017). We also found a negative correlation between social interaction and fentanyl preference, similar to the morphine preference findings of (Cooper et al., 2017). Importantly, our mice were pair-housed throughout the fentanyl procedure. Previously (Engeln et al., 2021), we showed that when mice are single-housed, CSDS-susceptibility is associated with increased early cocaine self-administration. When pair-housed, CSDS instead decreases cocaine self-administration, analogous to increased anhedonia. Consistent with our cocaine work, (Alexander et al., 1978) showed socially isolated rats consume more morphine relative to socially housed rats. Thus, the small effects we report here may reflect a kind of “social buffering” that decreases opioid consumption, similar to the protective effects of social interaction on heroin and methamphetamine craving (Venniro et al., 2018, 2019). Future work should investigate how stress interacts with social housing conditions to sensitize or blunt opioid consumption.

Here we found only stress-susceptible female mice maintained increased preference by the last choice period. In humans, sex-differences in OUD are largely dependent on the opioid (Back et al., 2010; Lee and Ho, 2013). Recent evidence suggests that women have higher rates of prescription opioid use (Serdarevic et al., 2017) and are more likely to report using opioids to cope with negative affect relative to men (McHugh et al., 2013). Women also exhibit increased susceptibility to stress-related disorders (e.g. PTSD, depression) (Bangasser et al., 2019). While we do not see striking sex-differences in our behavioral readout of susceptibility in mice, the persistently increased opioid consumption in our female mice mirrors that of humans and of other recent mouse work (Monroe and Radke, 2020). When we compare mice from our modified forced/choice model (Shaham et al., 1992) to mice that had free-choice throughout the entire procedure, stressed males do not appear different from unstressed males. It is tempting to speculate that in the forced/choice cohort, stressed male mice exhibit increased consumption due to either tolerance that develops from the forced epoch, or a desire to relieve the negative affect caused by abstinence/withdrawal (Koob, 2020). Female mice reliably consume more fentanyl than male mice, even when provided free choice throughout. This is consistent with previously reported sex differences in oral opioid consumption (Alexander et al., 1978).

Social withdrawal and other anhedonia-like behaviors are commonly exhibited following opioid abstinence and withdrawal (Bai et al., 2014; Becker et al., 2017; Bravo et al., 2020). In the **forced/choice cohort**, all unstressed and previously resilient mice exhibited decreased social interaction; in the **choice cohort**, only previously resilient mice decreased social interaction. Given that **choice cohort** unstressed mice consumed less fentanyl as compared with **forced/choice** unstressed mice, this suggests the social-withdrawal is dose-dependent. Opioid abstinence in previously resilient **choice cohort** mice—despite the overall dose being smaller--may have been sufficient to tip them towards susceptibility. Regardless of forced/choice or choice, the CSDS-susceptible mice maintain their previous social interaction behavior, which we believe may reflect a “floor effect.” Future work examining severity of withdrawal symptoms or other stress-like behaviors could address if susceptible mice exposed to fentanyl are more or less stressed than their resilient drug-exposed counterparts.

Given that NAc structural plasticity is an important determinant of synaptic strength, and heavily involved in both stress and drug-related behaviors, we examined Rho GTPase expression. Interestingly, while female mice displayed the more persistent elevation in fentanyl preference after stress, we only found differential expression in stressed male mice. Further, the changes to gene expression were consistent in susceptible mice regardless of physical or vicarious stress. We found decreased expression of Rac1, RhoA, and Cdc42 in NAc of stress and **force/choice** male mice. This is largely consistent with decreased Rac1 expression in NAc after CSDS (Golden et al., 2013), and decreased active Cdc42 in mPFC after chronic unpredictable stress (Luo et al., 2020). Both acute morphine withdrawal (Cahill et al., 2018) and CSDS (Francis et al., 2017; Fox et al., 2020a) engage NAc RhoA signaling. Thus, the downregulated RhoA expression seen here may reflect a compensatory mechanism in males that help protects against elevated late fentanyl preference. In agreement with compensatory downregulation, Rock1, a downstream effector of RhoA, is decreased in striatum after protracted morphine withdrawal (Drastichova et al., 2021). However our data are difficult to interpret given the competing effects of fentanyl and stress. Additionally, RhoA expression and dendritic remodeling after CSDS are cell subtype specific (Fox et al., 2020b, 2020a), and there is no work on NAc GTPases after CWDS. Future work can dissect precise cell-type specific adaptations that may explain the effects of CSDS/CWDS on subsequent fentanyl consumption.

In summary, we show stress-susceptibility is associated with increased fentanyl consumption that presents differently in male and female mice. We also found sex differences in the expression of dendritic-complexity molecules in the nucleus accumbens—a region important for both stress and drug-related behaviors. Our findings here, together with our updated mouse model, will provide the basis for more precise investigations on mechanisms of stress-sensitized opioid use in both sexes of transgenic mice.

## Funding and acknowledgements

The authors wish to acknowledge Symphanie Key for technical assistance. This study was funded by R01MH106500, R01DA047943, and R01DA38613 to MKL, T32NS063391 to DF and NIH K99/R00 DA050575 to MEF.

## Notes

### Competing Interest Statement

The authors have declared no competing interest.

