## Supplementary material for "Chronic Physical and Vicarious Psychosocial Stress Alter Fentanyl Consumption and Nucleus Accumbens Rho GTPases in Male and Female C57BL/6 Mice": Table 1

| **Limk1-F** | 5’-TGG GCT AGA AGG CAG CTT TA-3’ |
| --- | --- |
| **Limk1-R** | 5’- GGG ATT CAG ATC CCT GTC AA-3’ |
| **Rac1-F** | 5’- GCC ATG TAA CGC ACC TGT AA-3’ |
| **Rac1-R** | 5’- CAA AAG CTA GTC GGC TGG TC-3’ |
| **Rhoa-F** | 5’- GTG AAG CCT TGT GAA CGC A-3’ |
| **Rhoa-R** | 5’- TGA AAA GGC CAG TAA TCA TAC ACT-3’ |
| **Cdc42-F** | 5’- ACC TAC CCA CAT GCA CTC AT-3’ |
| **Cdc42-R** | 5’-ACT ATT ACT GGA AGG GCA AGG A-3’ |
| **Gapdh-F** | 5’- AGG TCG GTG TGA ACG GAT TTG-3’ |
| **Gapdh-R** | 5’-TGT AGA CCA TGT AGT TGA GGT CA-3’ |
